# The rs6505162 C>A polymorphism in the *miRNA-423* gene exhibits a protective element of coronary artery in a southern Chinese population with Kawasaki disease

**DOI:** 10.1101/596783

**Authors:** Jiawen Li, Jinxin Wang, Xiaoping Su, Zhiyong Jiang, Xing Rong, Xueping Gu, Huixian Qiu, Lanlan Zeng, Hao Zheng, Xiaoqiong Gu, Maoping Chu

## Abstract

**Background:** Manifesting as acute rash, fever and vasculitis, belonging to autoimmune syndrome, Kawasaki disease(KD) is prone to occur in infants and young children. Males and females is affected by KD at a ratio of 1.4 to 1.7: 1. KD is known to own many common clinical manifestations and complications, like coronary artery lesion(CAL) and coronary artery aneurysm(CAA). Polymorphisms of the rs6505162 locus in the *miRNA-423* gene are associated with enhancive susceptibility to coronary artery disease and the alterations of the four cytokines IL-4., IL-10, IL-21, IL-22 in the early stages of diabetes. However, no researcher has reported whether rs6505162 is related to KD susceptibility or no. Therefore, we carried out the trial concentrating on the connection between *miRNA-423* rs6505162 C>A polymorphism and KD susceptibility.

**Methods:** To obtain the genotypes of rs6505162 *in* objects enrolled by 532 KD children and 623 control, we applied Taqman real-time PCR and all statistical analyses was carried out by SAS.

**Results:** The comparison between all cases and all controls hinted that the rs6505162C>A polymorphism has no relationship with KD susceptibility. Nevertheless, a subgroup analysis revealed that the CA/AA genotypes of rs6505162 could reduce the occurrence of CAA (Adjusted age and gender odds ratio=1.30, 95%CI=1.02-1.67, *P*=0.037) and CAL (Adjusted OR=1.56, 95%CI=1.19-2.03, *P*=0.001)in KD patients.

**Conclusion:** Our final results stated clearly that *miRNA-423* rs6505162 polymorphism appears to be a protective element of CAL and CAA in southern Chinese suffers with KD.

## Introduction

In the year of 1967, the illness named Kawasaki disease(KD) was detected by Dr. Kawasaki.(1) Manifesting as acute rash, fever and vasculitis, belonging to autoimmune syndrome, Kawasaki disease(KD) which trigger acquired heart disease at most in non-developing countries is prone to occur in infants and young children.(2) In recent years, 18. 4% suffers are subjected to coronary artery disease and 20-25% untreated sick children suffered from coronary artery dilatation. The phenomenon above-mentioned could give rise to myocardial ischemia, and some of them develop into coronary aneurysm which perhaps rupture, even gigantic coronary aneurysms.(3, 4) KD can occur at any age including adults and neonate.(5–7) In patients with KD, complications of coronary artery disease which Initial corticosteroid therapy can prevent are related to the duration of disease before treatment.(8) The incidence of KD in most international areas such as Korea have been increasing slowly year by year, with a sex ratio of 1.4 to 1.7.(9–11)

The cause of KD is not totally clear and definite so far, and the exploration for the pathogen also has been upset.(12) According to the epidemiology and pathogenesis, the majority of human approve of the standpoint that the vasculitis of this illness is caused by the unsuitable immune reaction in individuals with hereditary susceptibility encountering one or more infectious irritants.(12, 13) In the culture of endothelial cells, somebody found that KLF4-miR-483 axis could be restrained by Kawasaki disease serum to speed up the development of Endothelial-to-Mesenchymal Transition(EndMT) which can injure vessel in KD patients.(14) Damage of endothelial cell homeostasis might concern the unusual circumstance of coronary artery in KD.(15) In a network analysis regarding protein interaction, the close contact between these genes concerning KD was verified to tell the pathogenesis of KD.(16) Several genes in the hypermethylated region were studied by Chen, and the correlation between the hypermethylated CpG locus and the pathogenesis of KD was mentioned for the first time. (17) These are some of the mechanisms of KD above.

Non-coding miRNAs, of which length are ~22 or so nucleotides, are produced by substances in the nucleus and cytoplasm and affect the course of genetics expression.(18) miRNA having a connection with plenty of mechanism and disease, include inflammatory response, severe asthma, diabetes mellitus, congenital heart disease, coronary artery disease(CHD). (19–23) Colorectal carcinoma whose potential biomarker might be *miR-423* rs6505162 has been studied by Jia, W.(24) Jha, Chandan K detected that The gene mutation in *microRNA-423* was in connection with enhancive susceptibility to CHD.(25)The *miRNA-423* rs6505162 having a certain relationship with CHD in prognosis, as studied in this article, is associated with HDL.(26) However, the connection of the *miRNA-423* rs6505162 C>A polymorphism with KD susceptibility has not been investigated so far. On the foundation of the medical center we employing resources from 532 KD patients and 623 controls implemented a new case-control study to appraise the relationship between the *miRNA-423* rs6505162 C>A polymorphism and the risk of KD in Han children from southern China at the point.

## Materials and methods

### Ethics declaration

The present study satisfying the standard of the Declaration of Helsinki was implemented in Guangzhou Women and Children’s Medical Center(2014073009), of which Review Committee acknowledged this study. And we ought to gather the written informed consent which legal guardians of participants endorsed.

### Study sample

A majority of participators coming from southern China have been recruited in the trial. All of participants who were from January 2012 - January 2017were Chinese Han with unrelated blood relationship. There were a plenty of samples that composed by 532 sufferers with recently diagnosed KD and 623 healthy controls in the research, according to the American Heart Association (AHA) reference.(27) Each participant offered 2ml fresh blood, of which 200ul was extracted for genomic DNA, and the specimens remained were stored for further study.

### Extraction and genotyping in DNA

On the basis of TIANGEN Company’s specification, genomic DNA was abstracted from 200ul blood of each participant by making use of TIANamp Blood DNA Kit (Centrifugal column, TIANGEN). Positive and negative samples were put into 384-well plates, which could be beneficial to contrast. Taqman real-time PCR was applied to genotype the *miRNA-423*rs6505162by ABI Q6 (Applied Biosystems).

### Statistical analysis in subgroups

In the group of the controls, genotype distributions that ought to be in line with the Hardy–Weinberg equilibrium were checked by a goodness-of-fit χ^2^ test. The diversity of selective variables were tested by utilizing two-sidedχ^2^ test as well as frequency distributions of the genotype. By means of univariate logistic regression, the relationship between the *miRNA-423* rs6505162 C>A polymorphism and the susceptibility of KD was described by odds ratios(ORs) and 95% confidence intervals (CIs). The adjustment of multivariate analysis was calculated through gender and age. Connections between susceptibility of KD and the genotypes were deeply assessed by stratification, when data were divided into subgroups about age, gender, coronary lesion and coronary artery aneurysm. The groups mentioned above are based on American Heart Association (AHA) and can be concretely grouped according to coronary artery and age.(27, 28)SAS software (version 9.4; SAS Institute, Cary, NC) in motion carried out all Statistical analyses quickly.

## Results

### Population feature

532 KD children and 623 healthy controls made up the participators in our research. The demographics entire participators possess were exhibited in the Table 1. 28.39 months was the average age of KD participators in onset. The quantity of 365(68.61%) in KD male patients was more than the one of 167(31.39%) in KD female patients. There was no distinct diversity in age (*P*=0.602) or gender (*P*=0.143) between KD children and healthy controls. There were 51(9.59%) patients with coronary artery aneurysm(CAA)when we took notice of the complications of KD. In addition, on the basis of coronary injury, 168(31.58%), 364(68.42%) cases were divided into CAL, NCAL, respectively.

**Table 1:**
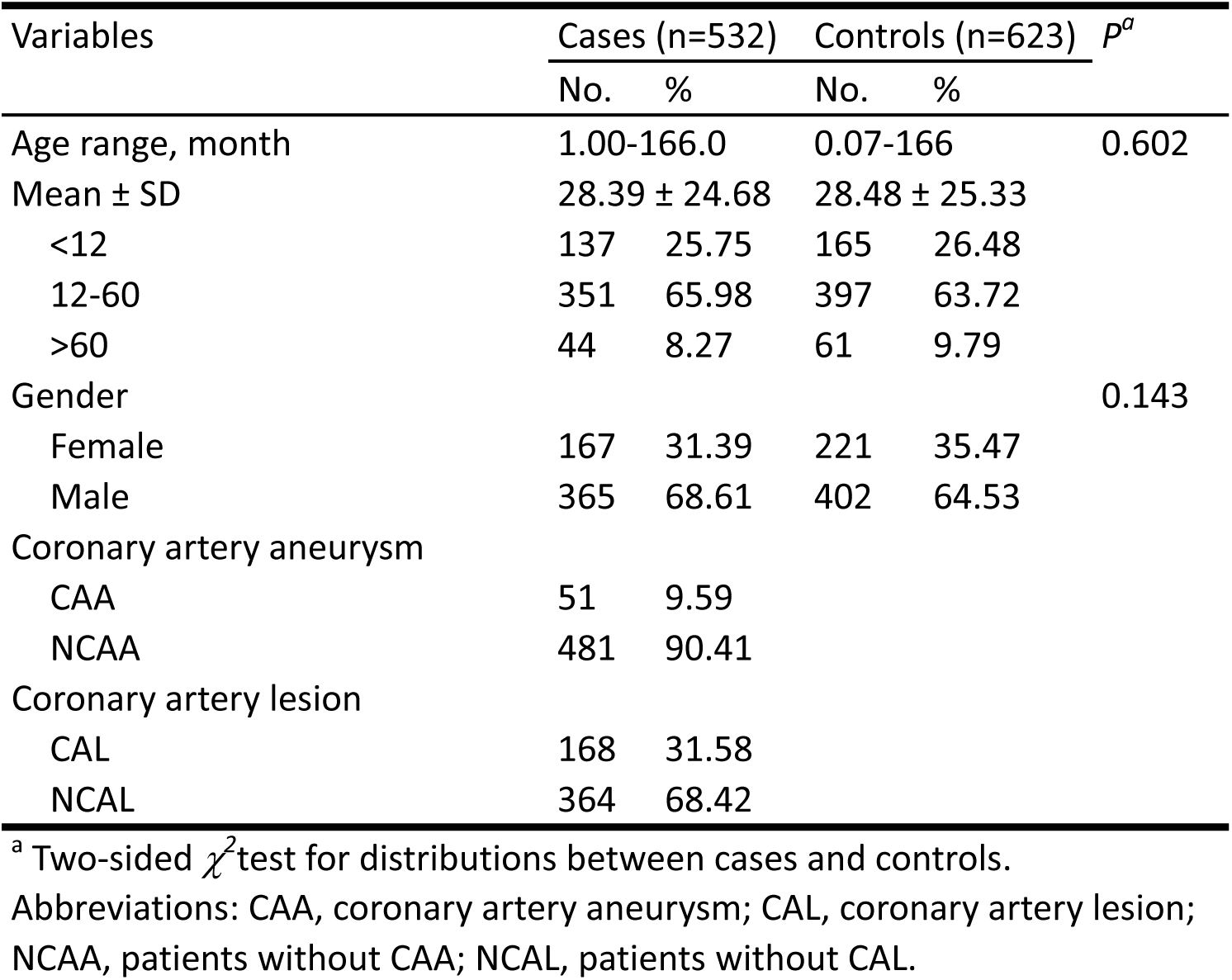
Frequency distribution of selected variables for cases and controls.

### Connection between the *miRNA-423* rs6505162 C>A polymorphism and KD susceptibility

In the groups of KD children and controls, to probe the association between *miRNA-423M* rs6505162 C>A polymorphism and KD susceptibility, we analyzed the genotype frequency distributions. As illustrated in Table 2, the controls satisfied the elements for Hardy-Weinberg equilibrium (P=0.791). The genotype frequency distributions of the *miRNA-423* rs6505162 C>A polymorphisms were 61.06% (CC), 34.97% (CA) and 3.97% (AA) in the KD group and 66.13% (CC), 30.50% (CA) and 3.37% (AA)in the controls. We observing the data from the rs6505162 C>A polymorphism and KD susceptibility, no significant connections was detected.

**Table 2.** Genotype distributions of rs6505162 C>A polymorphism and Kawasaki disease susceptibility

| **Genotype** | **Cases**<br>**(N=529)** | **Controls**<br>**(N=623)** | ***P*<sup>a</sup>** | **Crude OR**<br>**(95% CI)** | ***P*** | **Adjusted OR**<br>**(95% CI)<sup>b</sup>** | ***P*<sup>b</sup>** |
| --- | --- | --- | --- | --- | --- | --- | --- |
| CC | 323 (61.06) | 412 (66.13) |  | 1.00 |  | 1.00 |  |
| CA | 185 (34.97) | 190 (30.50) |  | 1.25 (0.98-1.61) | 0.075 | 1.25 (0.97-1.60) | 0.085 |
| AA | 21 (3.97) | 21 (3.37) |  | 1.29 (0.69-2.40) | 0.425 | 1.28 (0.69-2.38) | 0.443 |
| Additive |  |  | 0.202 | 1.20 (0.97-1.48) | 0.087 | 1.20 (0.98-1.48) | 0.084 |
| Dominant | 206 (38.94) | 211 (33.87) | 0.074 | 1.25 (0.98-1.59) | 0.074 | 1.25 (0.98-1.59) | 0.071 |
| Recessive | 508 (96.03) | 602 (96.63) | 0.688 | 1.19 (0.64-2.20) | 0.588 | 1.18 (0.64-2.20) | 0.591 |

<sup>a</sup> $\chi^2$ test for genotype distributions between Kawasaki disease patients and controls
<sup>b</sup> Adjusted for age and gender
Abbreviations: HWE, Hardy – Weinberg equation; KD, Kawasaki disease; OR, odds ratio.

### Stratification analysis in subgroups

Stratifying by age, gender, coronary injury(CAL), and coronary artery aneurysm (CAA)which are closely related to KD, we explored the connection between rs6505162 C>A polymorphism and KD susceptibility in a deeper level. In Table 3, there are two significant numbers we found. We noticed that the CA/AA genotypes of rs6505162 decreased the occurrence of CAA. (Adjusted OR=1.30, 95%CI=1.02-1.67, *P*=0.037). The CA/AA genotypes of rs6505162 could also decrease the occurrence of CAL(Adjusted OR=1.56, 95%CI=1.19-2.03, *P*=0.001). No other notable connections was detected, after we observed other subgroups such as gender and age.

**Table 3.** Stratification analysis for the association between rs6505162 C>A polymorphism and Kawasaki disease susceptibility

| Variables | CC<br>cases/controls | CA/AA | Crude OR<br>(95% CI) | *P* | Adjusted OR<sup>a</sup><br>(95% CI) | *P*<sup>a</sup> |
| --- | --- | --- | --- | --- | --- | --- |
| Age, month |  |  |  |  |  |  |
| <12 | 84/107 | 52/58 | 1.14 (0.71-1.83) | 0.580 | 1.11 (0.69-1.78) | 0.673 |
| 12-60 | 212/263 | 137/134 | 1.27 (0.94-1.71) | 0.119 | 1.28 (0.95-1.72) | 0.111 |
| >60 | 27/42 | 17/19 | 1.39 (0.62-3.14) | 0.426 | 1.32 (0.57-3.07) | 0.514 |
| Gender |  |  |  |  |  |  |
| Females | 96/148 | 71/73 | 1.50 (0.99-2.27) | 0.056 | 1.46 (0.96-2.22) | 0.081 |
| Males | 227/264 | 135/138 | 1.14 (0.85-1.53) | 0.393 | 1.13 (0.84-1.52) | 0.425 |
| Coronary artery aneurysm |  |  |  |  |  |  |
| CAA | 36/412 | 15/211 | 0.81 (0.44-1.52) | 0.518 | 0.82 (0.44-1.53) | 0.529 |
| NCAA | 287/412 | 191/211 | **1.30 (1.02-1.66)** | **0.038** | **1.30 (1.02-1.67)** | **0.037** |
| Coronary artery lesion |  |  |  |  |  |  |
| CAL | 122/412 | 46/211 | 0.74 (0.51-1.07) | 0.112 | 0.74 (0.50-1.07) | 0.112 |
| NCAL | 201/412 | 160/211 | **1.55 (1.19-2.03)** | **0.001** | **1.56 (1.19-2.03)** | **0.001** |

<sup>a</sup> Adjusted for age and gender. Statistically significant values are exhibited in bold ( $P < 0.05$ ).
Abbreviations: CAA, coronary artery aneurysm; CAL, coronary artery lesion; KD, Kawasaki disease; NCAA, patients without CAA; NCAL, patients without CAL; OR, odds ratio.

## Discussion

The nexus between KD susceptibility and the *miRNA-423* rs6505162 C>A polymorphism in our case-control investigation was further probed. No relevant association between the rs6505162 C>A polymorphism and KD susceptibility was noticed in the sick (Table 2). Using the subgroup analysis, we pointed out that the *miRNA-423* rs6505162 CA/AA genotypes is one of protective factors of CAA and CAL in KD patients (Table 3). These can serve as a basis for studying other relevant matters. Nevertheless, no conclusion that CAA and CAL in KD patients might have certain connection with age and gender has been analyzed in our subgroup research. More samples from KD patients with various age groups ought to be gathered to verify this discovery.

*miRNA-423* rs6505162 polymorphisms in KD children is first referred in this study. According to many reports, *miRNA-423* is closely related to heart failure and can also be used as a corresponding indicator, such as BNPs.(29–31) Other finding hinted that *miRNA-423* A allele and CA genotypes could enhance the risk of coronary artery disease(CAD).(25) Hakimzadeh reported that up-regulation of *miR423-5p* could identify the sick with low coronary collateral volume.(32) Alterations in the expression extent of 8 miRNAs including *miRNA-423* participating in the immune pathway might be involved in the immune process in the early stages of diabetes, through the alterations of the four cytokines IL-4., IL-10, IL-21, IL-22 and three pancreatic autoantibodies IA-2A, ICA, GAD65A.(33) In membranous glomerulonephropathy upregulation of *miRNA-423* had a certain significant association with down regulated IL6.(34)We found that the descending expression of the designated genes on proliferation and differentiation in myoblast was caused by the upregulated *miR-423-5p* suppressing the suppressor of fused homolog in expression.(35) In the polymorphism of *pre-MIR423* rs6505162, genetic mutations, C to A transition, inhibited the function of HEC-1b cell proliferation and migratory.(36) The abduction of EndMT progression accelerating cell proliferation and migratory capacity was caused by KD serum in endothelial cells, which could be one of the pathogenesis in KD.(14) On the basis of the series of articles above, there is a clear connection between *miRNA-423* and coronary artery and coronary collateral. At the same time, *miRNA-423* is associated with a range of inflammatory factors. Manifesting vasculitis, belonging to autoimmune syndrome, KD refers to inflammatory disease. Finally, the pathway of EndMT can be used to explain that *miRNA-423* can be a protective factor of KD as well as the stability of coronary arteries.

Our nowadays research suggested that the *miRNA-423* rs6505162 C>A polymorphism might not act on the susceptibility to KD in a majority of Han in southern China. KD is known to have many familiar clinical manifestations and complications, such as CAL and CAA.(3) We also noted that the CA/AA genotypes of rs6505162 reduced the occurrence of CAA and CAL in KD children. Nevertheless, our discovery including two positive point is required to affirm by more evidences owing to various factors restricting our study.

## Acknowledgements

Thanks to all the clinical samples belonging to Clinical Biological Resource Bank of Guangzhou Women and Children’s Medical Center, we can succeed in the trial. We thank An-qi Zhang for providing her technical guidance.

